# The influence of surfactin and glutamate on tomato root association by plant growth promoting rhizobacteria *Bacillus subtilis*

**DOI:** 10.1101/2025.04.10.647820

**Authors:** Rachel Warthen, Avaniek Cabales, Anna Wockenfuss, Aditya M. Kunjapur, Harsh Bais

**Affiliations:** Department of Plant and Soil Sciences, University of Delaware, Newark, DE, USA; Department of Chemical and Biomolecular Engineering, University of Delaware, Newark, DE, USA; AP Biopharma, University of Delaware, Newark, DE, USA

**Keywords:** PGPR, *Bacillus subtilis*, Tomato, roots, mature region, surfactin, glutamate

## Abstract

The influence of microbially and plant synthesized compounds on attachment of plant growth promoting rhizobacteria (PGPR) to various regions of the plant root are underexplored. Here, we examine the influence of surfactin and glutamate on the level and specificity of attachment of *Bacillus subtilis* along different regions of tomato root using chemical and genetic perturbations. First, we modify *Bacillus subtilis* PY79 to express a full-length phosphopantetheinyl transferase (*sfp*) native to other *B. subtilis* strains, and we observe surfactin biosynthesis under static incubation at 25°C in plant culturing media. We then perform microscopy using the wild-type PY79 strain, the sfp^+^ strain, and the wild isolate *B. subtilis* UD1022 to map the attachment of each strain along the entire root of young tomato plants incubated at 25°C, including when exogenously supplementing surfactin or glutamate. Through root mapping, we show that these strains prefer to attach near the mature region (MR) of the root and that colonization patterns vary based on exogenous metabolite concentration. Inclusion of glutamate in the media or through transient priming of the plant prior to bacterial inoculation strongly promoted root colonization by *B. subtilis* strains (both surfactin null and producers). In addition, the data shows that the domesticated strains were less efficient in binding compared to the wild *B. subtilis* strain. Interestingly, in the presence of glutamate, microbes lost their preference for association at the mature region, instead colonizing along the entire root. Overall, our work reveals a preference for association of these *B. subtilis* strains to the mature region of tomato in the absence of glutamate supplementation, confirms a strong influence of glutamate on root association, and demonstrates a smaller than anticipated role of biosynthesized or supplemented surfactin on root association, at least in a hydroponic culturing format.

## Introduction

Plant growth promoting rhizobacteria (PGPR) are known to induce both growth promotion and disease suppression in plants against classical bacterial and fungal pathogens. Among PGPRs, several *Bacillus sp*. strains are used as biocontrols for various diseases that impact crops (1). The majority of studied *Bacillus subtilis* are known for their PGPR traits that impact both plant growth promotion and triggers disease suppression in multiple plant species (2). The interaction of *B. subtilis* with plants is driving novel areas of research that seek to understand how benign microbes interact with plants for plant benefits. The impact of fundamental research on *B. subtilis*-plant interactions has led to commercialization of *B. subtilis* inoculums for sustainable agricultural practices (3). At this juncture, the knowledge to separate the fitness and biofilms formation ability of domesticated and wild strains *B. subtilis* is clearly understood (4). Lab strains of *B. subtilis* often show poor ability to colonize both biotic and abiotic surfaces suggesting lack of key microbial components for association (5). It is argued that nutrient rich media can facilitate the ability of some 168 derived strains to form effective biofilms (6). On the contrary, wild strains of *B. subtilis* shows robust colonization and biofilm formation ability on both abiotic and biotic surfaces.

While it is known that non-ribosomally synthesized peptides (NRPs) produced by *Bacillus sp*. exhibit antimicrobial properties, their effect on biofilm formation and root association is less well understood. Most of the antibiotics produced by *Bacillus* sps. belong to the class of non-ribosomal peptides (NRP) (7). The broad antimicrobial properties of NRPs are well documented, with their amphiphilic structure, NRPs such as surfactin, iturin, and fengycin/plipastatin can disrupt cell membranes and cause cell death (7, 8). Among bacterially derived NRPs, surfactin is a cyclic lipopeptide is known to be important in its role for antimicrobial properties (8). Surfactin production in *B. subtilis* strains is reregulated based on its interaction with the environment (9). A few strains of *Bacillus* sps. produce increased levels of surfactin against few plants and soil bacterial and fungal pathogens (10). The regulation and conditional production of surfactin synthesis and its impact on colonization on both biotic and abiotic surfaces is not known. Studies show that the surfactin biosynthetic mutants are equally capable of colonizing plants and abiotic surfaces (11). The differential expression of surfactin production in Bacillus spp. needs to be evaluated for its specific association with plant roots.

The ability of *Bacillus* spp. to colonize plant roots is dependent on *Bacillus* spp ability to form microcolonies and establish robust biofilms (12). It is shown that various plant derived/secreted molecules trigger biofilm operon in *Bacillus* spp. important for root colonization (10, 13). It was documented that a root secreted malic acid (MA) from plants under stress facilitates *Bacillus* spp. biofilm operons for root colonization (14). Likewise, other plant surface metabolites including plant polysaccharides are shown to trigger biofilm formation on the root surface and surfactin is not required for this association (13). However, possible mechanisms and root specific regional preference by PGPRs including *Bacillus* spp have not yet been disentangled. At this juncture, the root region-specific binding and its dependence on PGPR-mediated metabolites(s) including surfactin is not known.

Amino acids are known to induce defense responses and have been used as a fertilizer (15, 16). Among amino acids, L-Glutamate (Glu) administration has been known to modulate plant-microbe interactions; for example, Glu administration in rice plants develops resistance against *Magnoaporthe oryzae* (17) and the necrotrophic fungus *Alternaria alternata* in tomato fruit (18). It was further shown that exogenous supplementation of Glu in rice triggered pathogen resistance inducer in a salicylic acid (SA) inducible pathway (17). The role of Glu in Ca^2+^ transient response for plant insect interaction has been established (19). A recent study showed that Glu also works as a wounding signal that triggers long-distance defense signaling in a manner dependent on Glutamate like receptor (GLR) GLR3.3 and GLR3.6 (20). The role of Glu in plant-benign microbe interaction has not been established. Interestingly, supplementation of Glu in the media conditions is known to trigger the surfactin production of *Bacillus majovensis* A21 and *B. subtilis* 168 (21, 22). How supplementation of carbon sources such as Glu may interplay with microbial-derived metabolite such as surfactin inflicting plant colonization is not established.

To further investigate the mechanisms of Glu-surfactin interplay for root colonization in surfactin null *B. subtilis* strain (PY79), we created a root map in tomato plants under differential surfactin and exogenous Glu treatments. We show that the surfactin null strain PY79 failed to colonize roots of tomato plants. Exogenous surfactin treatment in the surfactin null strain PY79 reverted the colonization patterns specific to the mature region (MR) of roots. Interestingly, and contrary to literature reporting, the colonization observed in the domesticated *B. subtilis* strain with added surfactin was specific to the mature region (MR) of roots. Exogenous Glu treatments to the PY79 surfactin null or surfactin producing PY79 lines revealed an all-root colonization pattern compared to surfactin triggered MR specific colonization. In addition, exogenous surfactin and Glu treatment together showed efficient pellicle formation *B. subtilis* surfactin null and overproducing lines. Plant phenotype data showed plants treated with the PY79 or sfp^+^ mutant (with/without Glu) at different inoculation density didn’t modify the root phenotype except for length of lateral roots. Our results under hydroponic conditions tested suggest that Glu is an effective root-colonizer inducer (RCI) that overrides surfactin-dependent root colonization in tomato.

## Materials and Methods

### Seedling Preparation

Amish Paste tomato seeds (*Lycopersicon esculentum*) were purchased from Johnny’s Select Seeds. Seeds were soaked in water for at least 20 minutes and then suspended and shaken in 2:1 bleach for with a drop of TWEEN-20 for 20 minutes. Seeds were rinsed 3 times with sterile water and then shaken 70% ethanol (EtOH) for 30 seconds. Seeds were placed on ½ MS (Murashige and Skoog, 1962). MS agar plates under photoperiod of 12-hour day 12-hour night cycle for 7 days at a photosynthetically active radiation (PAR) of ∼100 µmol m^-2^ s^-1^.

### Culturing Bacteria

*Bacillus subtilis* strains (including UD1022) were cultured in LB media at incubated at 37℃. Antibiotic selection was used for selection with certain strains. For PY79::mNeongreen, kanamycin was used at a concentration of 5 µg mL^-1^, and for the PY79 surfactin producing strain (PY79 lacA::Pveg-sfp::cat; from here on, PY79 sfp^+^) chloramphenicol was used at a concentration of 5 µg mL^-1^. For plant incubation experiments, strains grown overnight in liquid LB with appropriate antibiotic concentration was used (See Table 1).

**Table 1:** List of bacterial strains used in the study.

| Bacterial strain | Antibiotic Selection | Reference |
| --- | --- | --- |
| *B. subtilis* PY79 | No selection | ATCC |
| *B. subtilis* PY79::mNeongreen | Kanamycin 5 μg mL<sup>-1</sup> | Gift from Devon Stork, Ethan Garner, and George Church |
| *B. subtilis* PY79 lacA::Pveg-sfp::cat (sfp<sup>+</sup>) | Chloramphenicol 5 μg mL<sup>-1</sup> | This study |
| *B. subtilis* UD1022 | No Selection | Bishnoi et al., (2015) |

### Surfactin analysis and creation of surfactin overproducing strain. Surfactin Quantification

Surfactin quantification was done via high performance liquid chromatography with an isocratic method of acetonitrile with 1% trifluoroacetic acid. 50 μL of sample was injected and run for 10 minutes, and absorbance was analyzed at 205 nm. Quantification was accomplished by comparing peak areas of three major peaks in the commercial standard. Surfactin was purchased from Smolecule (San Antonio, TX, USA). Stocks were made by dissolving surfactin in deionized water and diluted as needed.

### Strain Generation

Polymerase chain reaction (PCR) based DNA amplification was performed with KOD Xtreme Hot Start Polymerase. PCR primers were designed to have 20 bp of overlap for ligation via Gibson isothermal assembly. The assembled construct was PCR amplified before purification and then used for transformation. The gene encoding Sfp was purchased from the Bacillus Genus Stock Center (BGSCID: ECE221).

Transformation was done by inoculating 1 mL of MC media from plates. At 4 h, the DNA was added to 200 µL cells and grown at 37°C for 2 more hours before plating on selective media. MC media was made at a 10× stock. The 10× stock has final concentrations of 1 M potassium phosphate pH 7, 30 mM sodium citrate, 20% (w/w) glucose, 220 mg/mL ferric ammonium citrate, 1% casein hydrolysate, 2% potassium glutamate. Aliquots were stored at −20C. MC media was made for the transformation and supplemented with 3mM MgSO_4_. Colonies were verified to have the DNA by colony PCR, where cells were suspended in 50 μL TE +10μm glass beads, then vortexed for 10 min, boiled for 30 min, vortexed for 10 min. 1 μL was used as a PCR template. Purified colony PCR DNA was sequenced by Plasmidsaurus to confirm. PCR primers were purchased from Integrated DNA technologies (IDT) (Coralville, IA). KOD Xtreme Hot Start and trifluoroacetic acid was purchased from MilliporeSigma (Burlington, MA, USA). Acetonitrile was purchased from RICCA (Arlington, TX, USA).

### Inoculation for Root Maps

Seven-day old tomato plants were taken from ½ MS plates and suspended in 1.9 mL of liquid media in 12-well plates. *Bacillus subtilis* strains grown in liquid with appropriate antibiotic selection were pelleted and resuspended in liquid media ½ MS and diluted from the initial stock to 10^7^ cells mL^-1^ (OD^600^ = 0.20). Plants grown in liquid ½ MS with 15 g L^-1^ sucrose were treated with the ∼10^6^ cells mL^-1^. Inoculated plants were incubated at a photosynthetically active radiation (PAR) of ∼100 µmol m^-2^ s^-1^ with a 12:12 hr photoperiod. Plants were monitored for root mapping at 72 h post treatment.

### Staining

Non-reporter strains such as PY79 surfactin producing strain (PY79 sfp^+^) and *B. subtilis* UD1022 (undomesticated strain) were stained with calcofluor white (dilution 1:1000) for 15 minutes and then with SYTO13 (dilution 1:1000) for 30 seconds before being mounted for bioimaging. SYTO13 channels overlaying brightfield channels were utilized to assist images containing non-reporter strains (PY79 sfp^+^ and UD1022) in appearing more uniform to images containing reporter strains such as PY79::mNeongreen constructs.

### Plant phenotyping

Tomato (Amish Paste tomato) plants were germinated and maintained in ½ MS media. Plants post germination (7 days old) were co-inoculated with different inoculum densities (OD_600_ = 0.0001, 0.001, and 0.01) with either PY79 or PY79 sfp^+^ strains. Plants inoculated with either PY79 or PY79 sfp^+^ strains were grown in the presence, or absence of L-glutamate (5 mM) in MS media. Root phenotypes such as length of primary roots (PR), length of lateral roots (LLR) and number of lateral roots (NLR) were quantified after 7 days of incubation.

### Microscopy

Roots were imaged from the first lateral root down to the root tip using the Stellaris 8 tauSTED/FLIM Confocal Microscope. The LasX program was used to input scale bars and prepare images to be made into figures.

### Root Measurements

Root sections were measured using ImageJ software and lif files of merged regions.

### Pellicle Assay

For the pellicle assays, the overnight grown and plated bacterial cells were selected from a plate and placed in 50 mL flacon tubes containing 10 mL of LB broth and appropriate antibiotics. Liquid cultures were left in 37℃ for 24 h, cultures were removed from the shaker and diluted to an OD600 of 0.2. Cultures were then placed back in the shaker. The cultures were further diluted in 1 mL of MSgg media, the pellet culture assays were set in 24-well plates. Plates were further incubated at the room temperature or placed in 30℃. Pellicles were harvested and photographed at 72 h post inoculation.

### Data analysis and statistics

Primary root length, number and length of lateral roots were measured using ImageJ. A one-way ANOVA was performed to determine the effects of differing bacterial concentrations (SOM Figure 1), as well as a two-way ANOVA to observe the effect of two bacteria (one concentration), in the absence or presence of glutamate (SOM Figure 2). A Tukey multiple comparison was calculated from the means using GraphPad Prism, at a significance level of p < 0.05.

**Figure 1:**
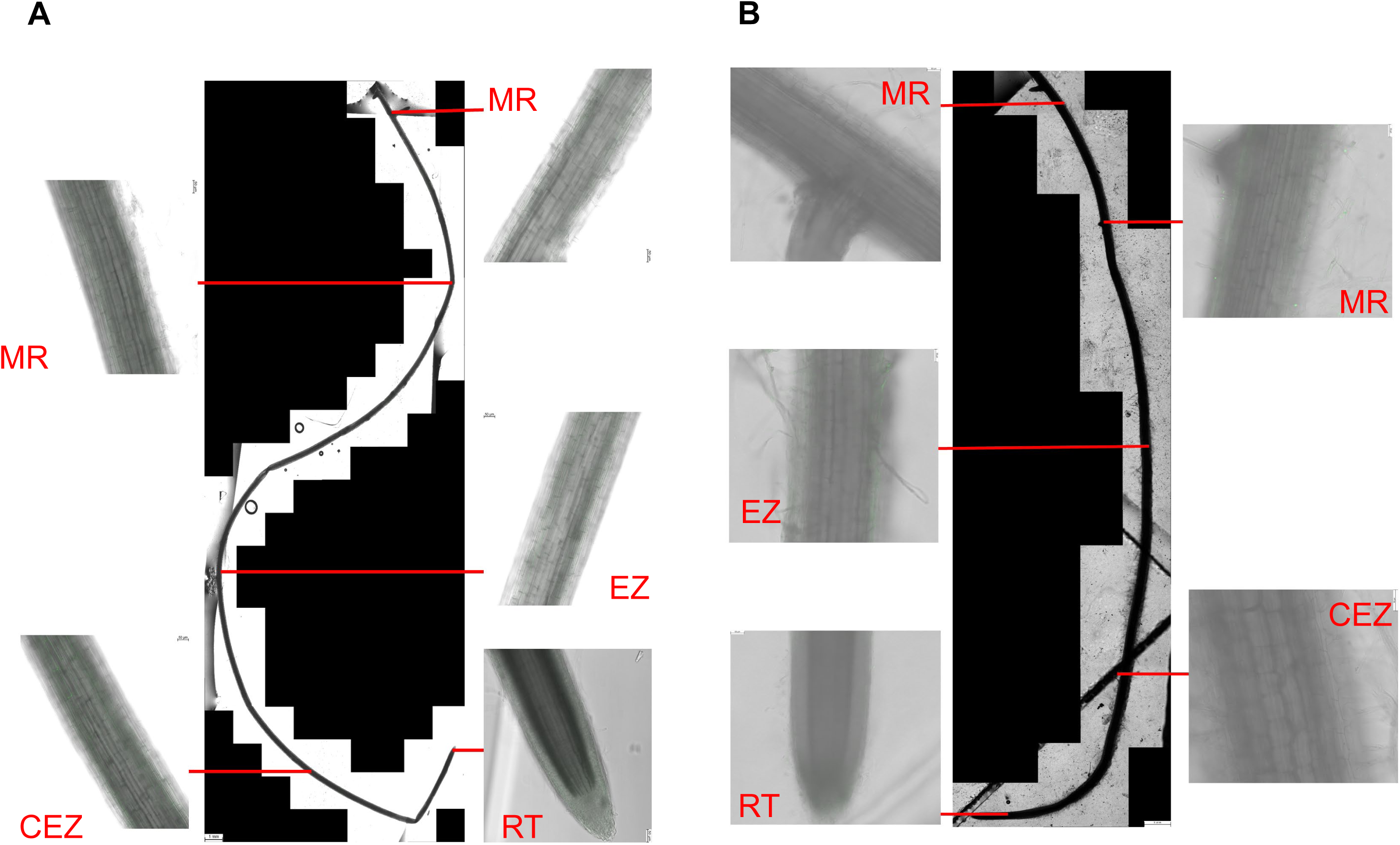
Root attachment of a lab strain of *B. subtilis* (PY79) (sfp-) on tomato plants. Plants were imaged 72-hrs post inoculation. *B. subtilis* PY79::mNeongreen (sfp-) was added at 10^7^ cells ml^-1^. The panel depicts root maps of an uninoculated root (**A**) and *B. subtilis* PY79::mNeongreen (**B**). Root regions are indicated by either MR (Mature Region), EZ (Elongation Zone), CEZ (Central Zone of Elongation, or RT (Root Tip). Scale bar= 5mm.

**Figure 2:**
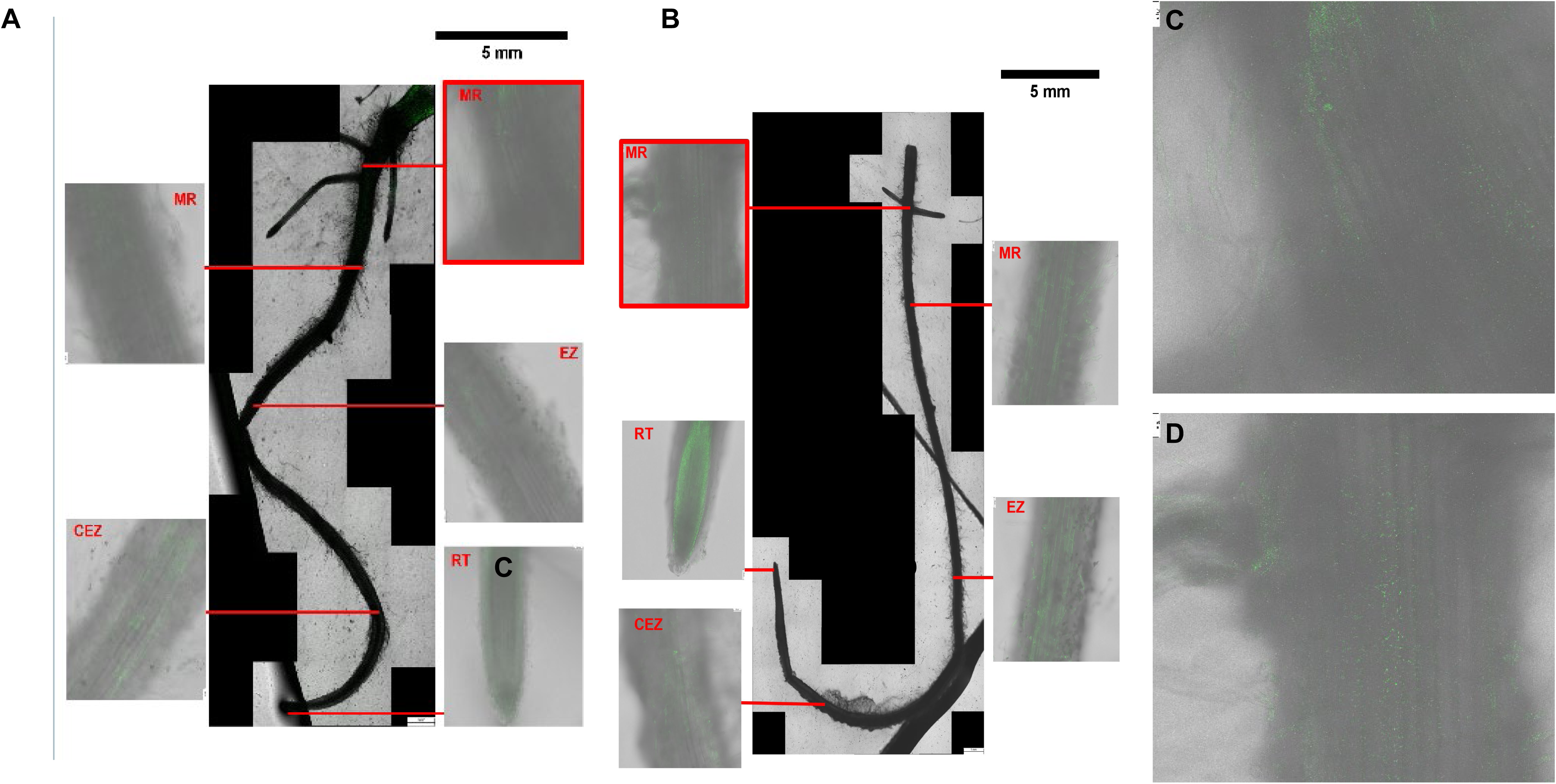
Exogenously supplemented surfactin and its role in root association. Plants were imaged 72-hrs post inoculation. Prior to inoculation plants were primed with *B. subtilis* PY79::mNeongreen was added at 10^7^ cells ml^-1^ for 48-hrs with surfactin (A and B). Samples were also either washed (A) or left unwashed (B). Panels (C) and (D) are enlarged regions from the mature region from surfactin washed and unwashed treatments. Red boxes indicate areas in which colonization was observed. Root regions are indicated by either MR (Mature Region), EZ (Elongation Zone), CEZ (Central Zone of Elongation, or RT (Root Tip). Scale bar= 5mm.

## Results

### Root attachment of a lab strain of *B. subtilis* (PY79)

To evaluate the specific root colonization patterns by different *B. subtilis strains*, we created a root map in young tomato seedlings. Tomato seedlings (7 day old) grown in vitro were inoculated with various bacteria and different regions of roots were imaged finely for root specific colonization. The entire roots were imaged starting from root tip to the first lateral position post treatment. Tomato roots inoculated with the surfactin nonproducing PY79 strain showed no colonization in any root region 72 h post-inoculation (Figure 1). The non-surfactin producing strain PY79 displays no root attachment and appears like the untreated control plant which had not been treated with any benign strains of bacteria (Fig. 1A-B). While it is known that PY79 is deficient in biofilm formation, its ability to bind tomato root has not been investigated, and some studies have suggested that it and other *B. subtilis* strains might have this ability (23). We demonstrate that, under conditions tested, it is unable to associate with tomato roots.

### Exogenously supplemented surfactin and its role in root the association

To observe its potential role in root association, surfactin (25 µg ml^-1^) was exogenously added to the minimal media to prime plants for 48 h prior to the addition of surfactin null strain PY79 (srf-). Surfactin was then either left in the media (unwashed) or plants were moved to new fresh media not containing surfactin (washed) at the time of inoculation. Minimal attachment was observed 72 h post-inoculation on both washed and unwashed plants by PY79 in the mature region, with unwashed plants having slightly more colonization than washed plants. Colonization was observed on the mature regions (MR) of the root and only in these regions. Both the unwashed (Figure 2) and washed (Figure 2) conditions contain colonization in the mature regions of the root. The data suggests that the exogenously surfactin supplemented roots facilitate subtle root association by a surfactin null strain PY79. In addition, the preferential binding by surfactin null strain PY79 in the MR region of the root suggests an unknown conducive signaling favoring MR region association by the bacteria.

### A surfactin overproducing strain is not a successful colonizer

Having shown that an exogenous supplementation of surfactin to plants with a surfactin null strain PY79 (sfp-) revealed a subtle colonization in the MR region of roots, we wondered whether a strain capable of surfactin biosynthesis may also exhibit a similar phenotype. To this end, a surfactin overproducing strain (PY79 sfp^+^) was used to inoculate roots and colonization data was imaged post 72 h of inoculation. We engineered a surfactin producing PY79 (sfp+) by genome integration of the *sfp* gene under constitutive control by the P_veg_ promoter into the *lacA* locus. This PY79 sfp^+^ strain relies on the native *srf* operon and is under native regulation for the assembly of surfactin. Despite colonization being observed in Figure 2 where surfactin was added to the media, colonization was not observed with the surfactin overproducing PY79 sfp^+^ strain (Figure 3). A root map of the undomesticated *B. subtilis* strain UD1022 (UD1022) was also created to compare how this undomesticated strain colonized the roots compared to the domesticated strains. The undomesticated strain UD1022 colonizes both the mature region and in some cases the elongation zone (EZ) as well (Figure 3). The data suggests that surfactin treated PY79 showed subtle root colonization phenotype compared to a surfactin overproducing lines. In addition, the undomesticated *B. subtilis* strain UD1022 showed MR and EZ specific colonization compared to surfactin treated PY79.

**Figure 3:**
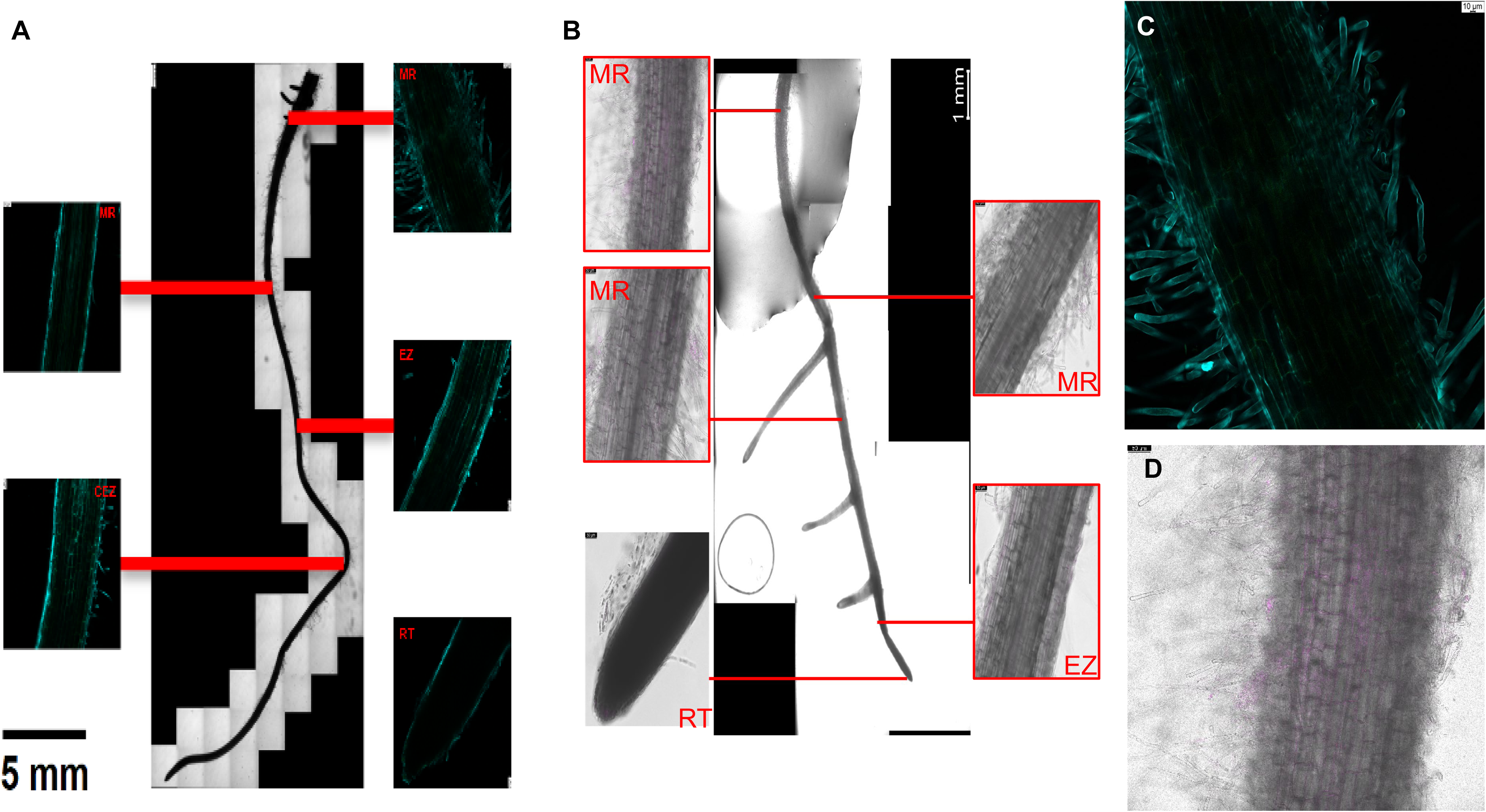
Root attachment of surfactin overproducing strain (sfp+) on tomato plants. Plants were imaged 72-hrs post inoculation. *B. subtilis* (sfp+) was added at 10^7^ cells/mL. Images of a surfactin overproducer sfp+ (A). Tomato roots were also inoculated with an undomesticated *B. subtilis* strain (UD1022) (B) for colonization. (C) and (D) are enlarged images from the mature region of the previous treatment of a sfp+ and UD1022 respectively. The roots and non-reporter strains were fixed and stained with calcofluor white and SYTO13. Root regions are indicated by either MR (Mature Region), EZ (Elongation Zone), CEZ (Central Zone of Elongation, or RT (Root Tip). Scale bar= 5mm.

### Surfactin analysis in the overproducing strain

We showed that the sfp^+^ strain failed to colonize the tomato roots suggesting the conditional specific response of surfactin in root colonization. To evaluate if the sfp^+^ strains were able to biosynthesize surfactin, we tested the growth and production of the sfp^+^ producer strain under static incubation at 25°C (Figure 4). The data showed that sfp^+^ strain as able to biosynthesize surfactin at 72 hrs of incubation, suggesting a density-derived surfactin biosynthesis (Fig. 4).

**Figure 4:**
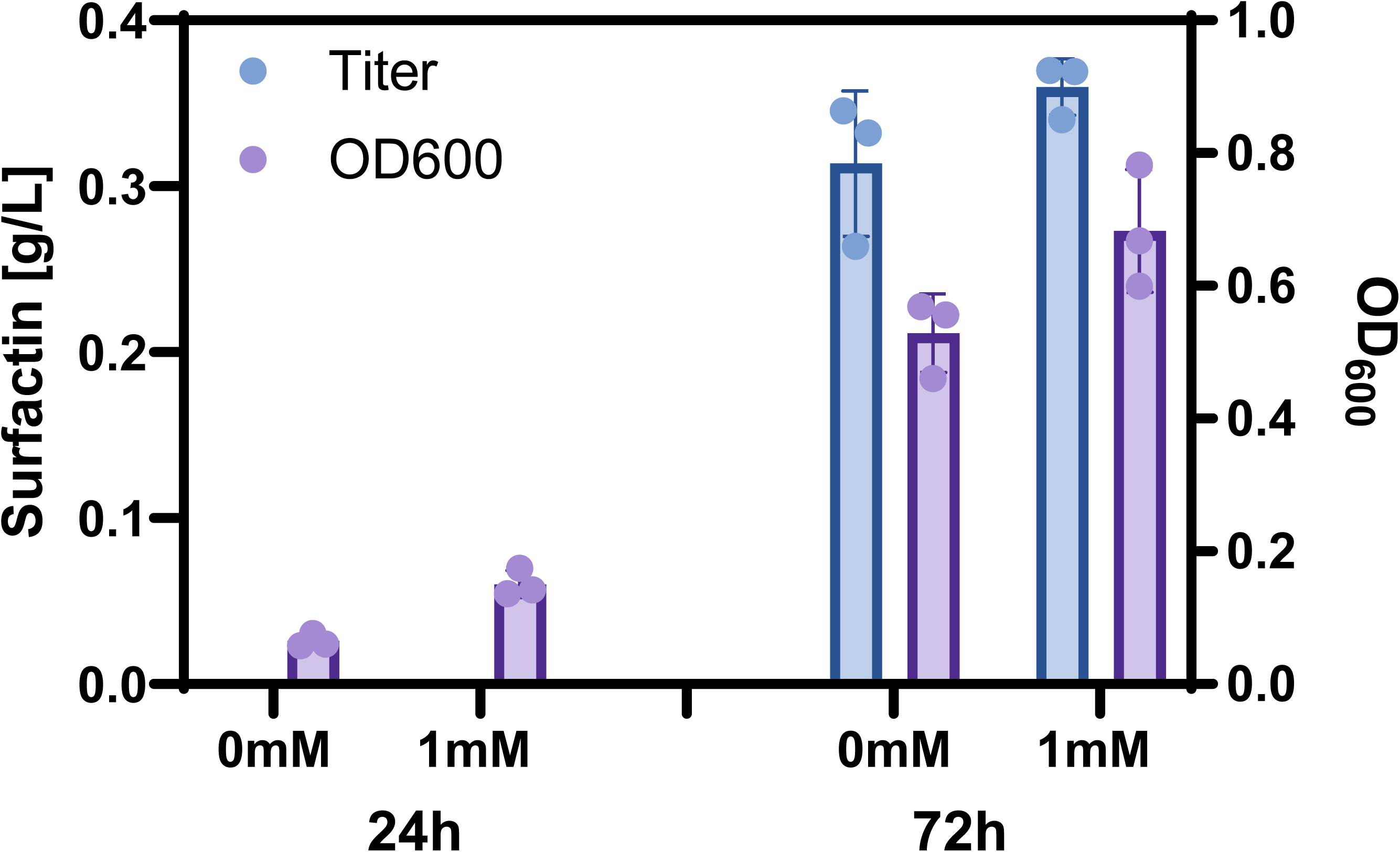
Surfactin production and cell growth of PY79 sfp+ under static conditions, incubated at 25°C. Surfactin titers and cell growth of sfp+ strain was monitored post 1mM glutamate supplementation under static condition. Both surfactin titer and growth (OD600) was measured 72 hr post glutamate treatment.

### Glutamate assists with the root attachment in PY79 species

To analyze the role of a known PRI L-Glutamate (hereafter: Glu) in surfactin interplay for root colonization phenotype tomato roots were treated with glutamate (5 mM). Exogenous treatment of Glu to tomato roots with surfactin null strain (PY79) showed enhanced root colonization in the MR region of the roots (Figure 5). Attachment was primarily observed within the mature region of the root in both PY79 and PY79 sfp^+^ and does not extend to either the zone of elongation or the root tip (Figure 4). A similar trend also appeared to occur with PY79 sfp^+^, wherein surfactin overproducing line showed root colonization in the MR region with exogenous treatment of Glu (5mM) (Figure 5). This data suggests that addition of Glu a known PRI may trigger root specific colonization in the MR region independent of surfactin.

**Figure 5:**
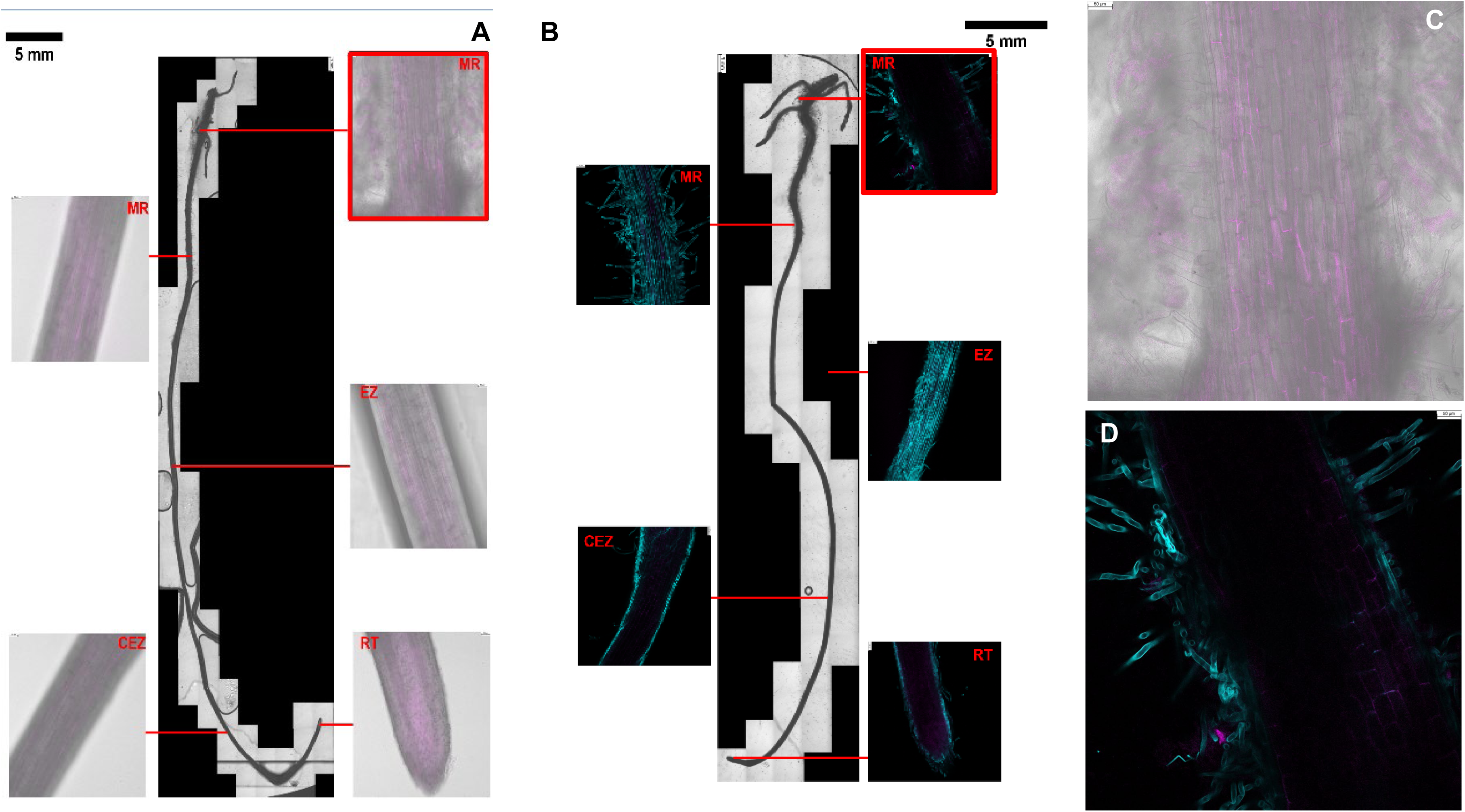
Impact of glutamate on root attachment in *B. subtilis* PY79::mNeongreen (sfp-) and sfp+ strains. Glutamate was added to media at a concentration of 5 mM. The entire root map (A-B) showed association of surfactin null strain and an overproducing line. The inset images (C-D) showed the colonization patten in the MR region of the roots. The roots and non-reporter strains were fixed and stained with calcofluor white and SYTO13. Red boxes indicate areas in which colonization was observed. Root regions are indicated by either MR (Mature Region), EZ (Elongation Zone), CEZ (Central Zone of Elongation, or RT (Root Tip). Scale bar= 5mm.

### Glutamate supplementation negates surfactin biosynthesis in static bacterial cultures

We observed Glu supplementation triggering root colonization in sfp^+^ strains, next we evaluated surfactin biosynthesis is favored under static culture conditions with or without supplementation of 1 mM L-Glu. Under these conditions, we observe comparable growth and surfactin production over the 72-hour period in both conditions (Figure 4). These data indicate that Glu supplementation may not directly interface with surfactin biosynthesis, at least under static culture conditions.

### Priming roots with Glu and surfactin trigger robust colonization by PY79

Having shown that plants treated with exogenous Glu and inoculated with surfactin null and overproducing line showed similar root colonization phenotype, we evaluated the role of exogenous surfactin in the Glu-surfactin interplay for the root colonization phenotype. When plants were primed with both Glu and surfactin prior to inoculating roots with surfactin null strain PY79, robust colonization was observed in multiple regions of the roots. This colonization occurs in both washed and unwashed conditions at a concentration of 5 mM of Glu. Addition of Glu (5 mM) did not alter the plant or microbial growth (data not shown). Interestingly, plant root cultures that were subjected to wash treatment showed less colonization than that of the unwashed Glu treatments (Figure 6). Contrary, to the surfactin treated roots and MR specific colonization, roots treated with Glu and surfactin together showed root colonization not specific to MR and colonized the entire roots (Figure 6). The phase image of the entire root map also showed the entire root colonization by PY79 surfactin null strain treated with exogenous surfactin and Glu (Figure 6). The data suggests that combination of surfactin and Glu may override root specific colonization patterns triggered by lone Glu/surfactin treatment.

**Figure 6:**
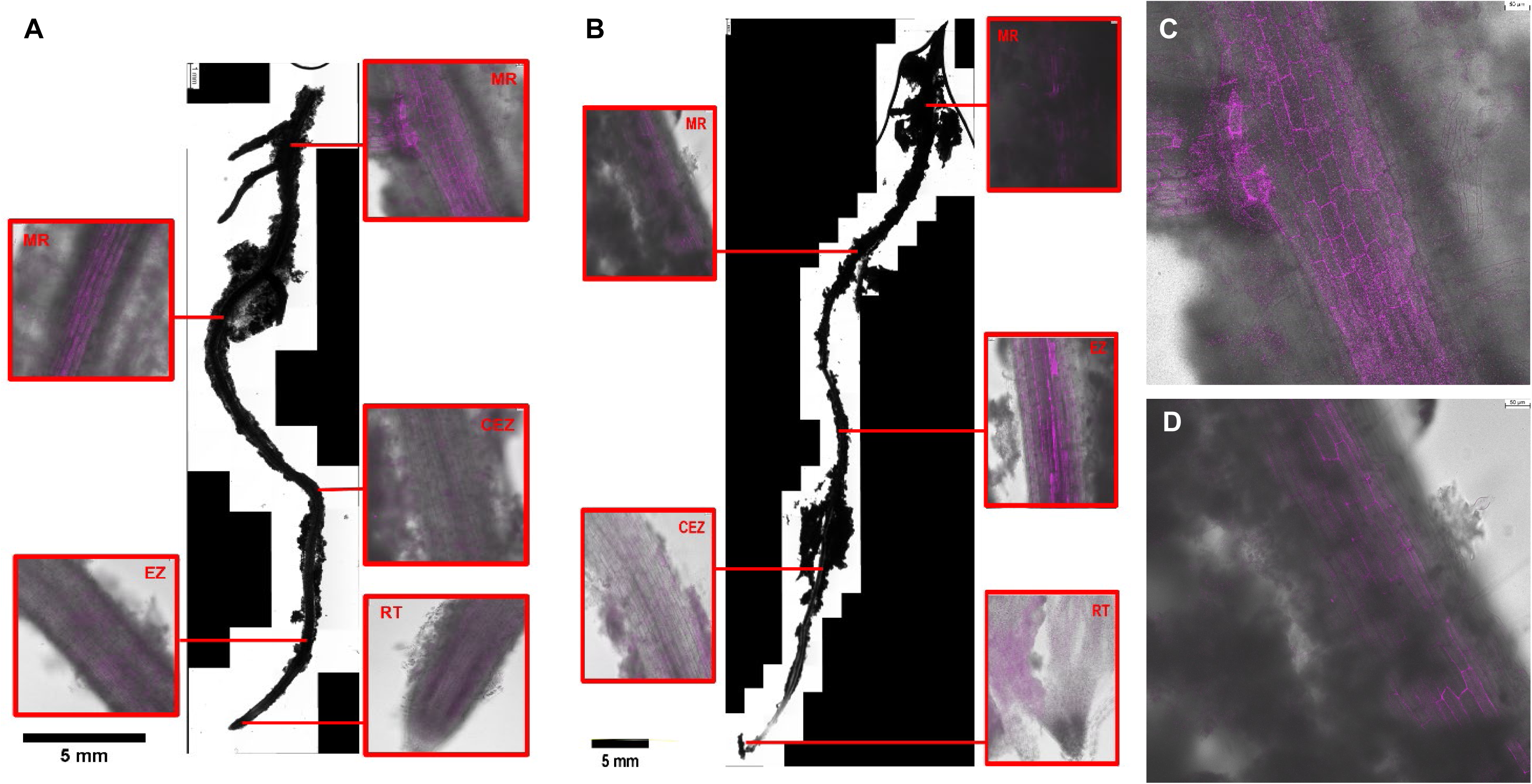
Impact of glutamate and surfactin priming (washed [A] and unwashed [B]) on root attachment in *B. subtilis* PY79::mNeongreen (sfp-). Glutamate was added to media at a concentration of 5 mM of glutamate alongside surfactin. Roots primed with surfactin were left washed or unwashed prior to glutamate supplementation for root mapping. The panel C and D shows the inset images of the MR root region. Red boxes indicate areas in which colonization was observed. Root regions are indicated by either MR (Mature Region), EZ (Elongation Zone), CEZ (Central Zone of Elongation, or RT (Root Tip). Scale bar= 5mm.

### Glutamate supplementation induces surfactin biosynthesis in plant-*B. subtilis* cultures

We tested the surfactin biosynthesis in tomato plants inoculated with surfactin overproducing strain (sfp^+^) supplemented with L-Glu (5 mM). Surfactin overproducing strain (sfp^+^) when cultured with the tomato plant and 5 mM glutamate and extracted for surfactin showed presence of surfactin as the standard surfactin HPLC traces (SOM Figure 1). As expected, neither the plant nor plants inoculated with PY79 (srf null strain) showed presence of surfactin peaks (SOM Figure 1). Contrary to the bacterial static culture surfactin analysis with L-Glu, the plant associated sfp^+^ strain with exogenous surfactin shows traces of surfactin suggesting that L-Glu may elicit surfactin biosynthesis in sfp^+^ strain.

### Glutamate aids in the development of thicker pellicles

Having shown that Glu and surfactin exogenous application to tomato roots changed the root colonization pattern, we evaluated the role of Glu in *B. subtilis’* ability to form pellicle on abiotic surfaces. In addition, pellicle experiments were conducted in parallel with the dual temperatures of 30℃ and 22℃ (room temperature) to observe if temperature had any impacts on pellicle formation. Abiotic surfaces appear to display different results from the biotic surfaces seen in the root maps, specifically in the cases of PY79 and sfp+ strain. Both surfactin overproducing line and undomesticated strain UD1022 formed robust pellicles independent of Glu/surfactin or temperature conditions (Figure 7). Interestingly, undomesticated strain UD1022 formed wrinkled pellicles under Glu treatments compared to the lab strains (Figure 7). There is also a pigment change in both sfp+ and UD1022 at 30°C (Figure 7). While this change only occurs in glutamate treatments for sfp+ strain, UD1022 appears to turn a darker yellow under at 30°C in all treatments. The data suggests pellicle formation in *B. subtilis* is independent of Glu and surfactin treatment compared to Glu dependent biotic surface colonization.

**Figure 7:**
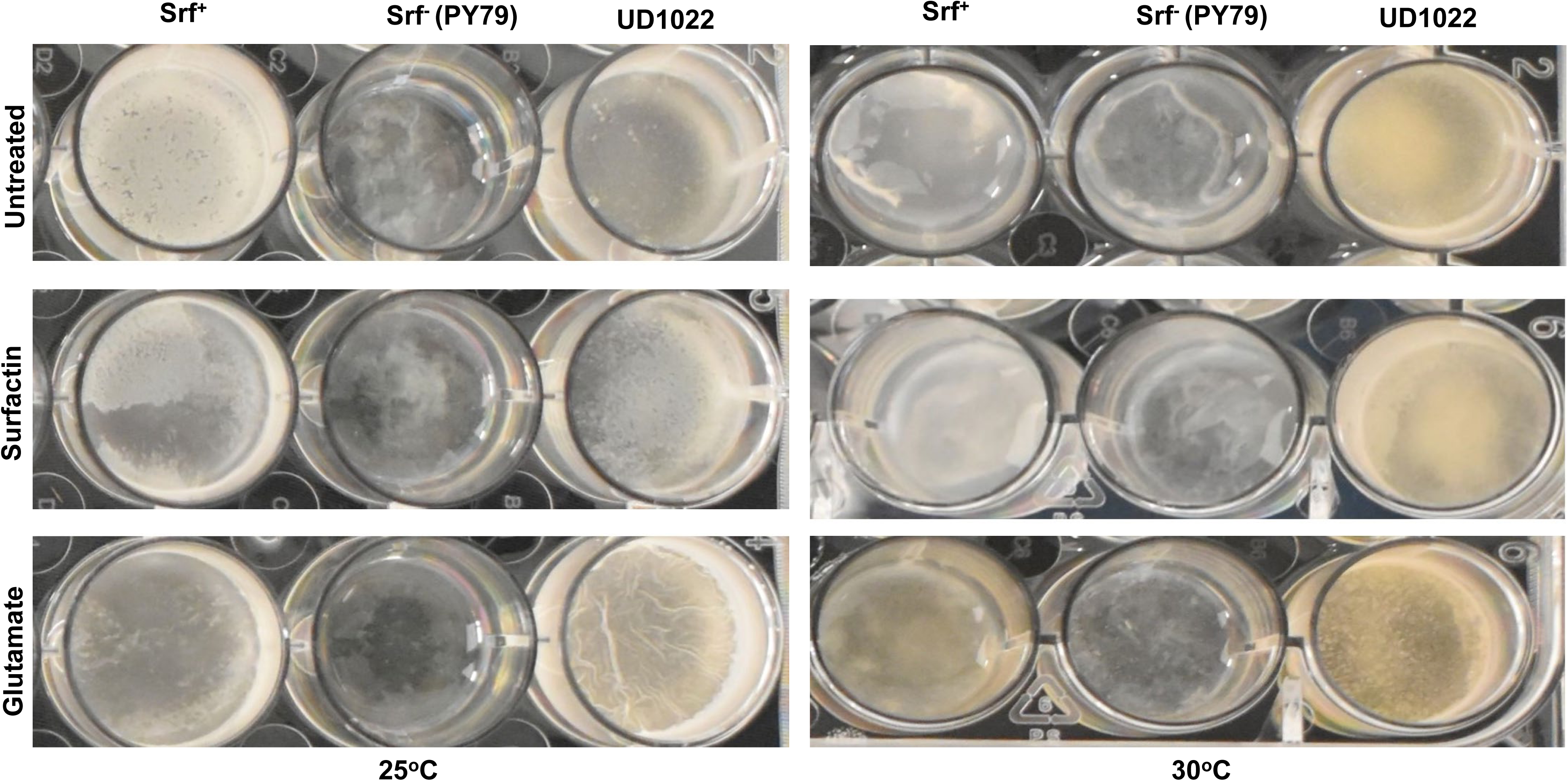
Effect of temperature, glutamate and surfactin on pellicle formation in lab and undomesticated *B. subtilis* strains. A) Pellicle formation of surfactin null strain (PY79) (sfp^-^), surfactin overproducer (sfp^+^) and a wild *B. subtilis* strain (UD1022) grown at 25°C. B) Pellicle formation of surfactin null strain (PY79) (sfp^-^), surfactin overproducer (sfp^+^) and a wild *B. subtilis* strain (UD1022) grown at 30°C. Different conditions including exogenous supplementation of surfactin (25 µg ml ^-1^), and glutamate (5 mM) were added to the media. Images of the pellicle were photographed on 72-hrs post inoculation.

### Role of surfactin and Glutamate on root phenotype

To analyze the role of surfactin and glutamate supplementation with tomato plants, plants were treated with PY79 or PY79 sfp^+^ and grown with different bacterial inoculum densities. Plants post incubation were analyzed for any root phenotype in terms of length of primary roots (PR), number of lateral roots (NLR) and length of lateral roots (LRR). Though the plants treated with higher inoculum of PY79 showed reduced LRR, they were not significantly different from lower inoculum densities (SOM Figure 2A-B). The sfp+ strain showed an inverse LRR phenotype in the lower density treatment with reduced LRR, compared to the higher inoculum, but was not statistically different than the other treatments (SOM Figure 2A-B). The data showed that supplementation of glutamate to tomato plants altered the colonization patterns which was srf independent. The root phenotyping data showed that there were no significant changes to the LPR, and NLR phenotype post glutamate supplementation. Surprisingly, plants exposed to glutamate showed a decrease in LLRs (SOM Figure 2). Plants treated with glutamate and srf + showed significantly increased LLR compared to the srf null strain (SOM Figure 3).

## Discussion

To protect crops from plant pathogens to reduce disease incidences, we need to better understand the role of PGPRs in the soil that interact with plants and plant-pathogenic microbe interactions. *Bacillus* species are commonly found in disease suppressive soils and is often used in biocontrol methods to help prevent disease in many crops (24, 1). *Bacillus subtilis* is often studied as it is known to not only to prevent disease but promote plant growth as well (2). The interaction of PGPRs and mainly *Bacillus* spp. with plants is mediated through root exudates (25). Though much attention has been focused on how root exudates influence soil and rhizosphere microbiome, the impact of root region specific exudations and its role in root colonization by PGPRs is not known. It is shown in many plant species that the root secretions modify rhizosphere colonization by PGPRs but how different regions of roots modulate secretion profile while interacting with rhizobacteria may shed more light on preferential colonization pattern of PGPRs.

Surfactin is known to regulate critical microbial functions including antibiosis against major bacterial species (26). The role of surfactin in modulating plant-microbe interactions has been questioned and debated. Some reports suggest that surfactin is an important metabolite that regulates root colonization and disease reduction against pathogens in plants (27). In contrast, Thérien et al. (28) reported that surfactin may not play a critical role in root association. The gaps in the literature also do not clarify the specific role of plants in mediating surfactin dependent colonization. For example, at this juncture we do not know which region of the root prefers PGPR colonization and if root-specific association of *Bacillus* sps is surfactin dependent. In addition, the *in vivo* surfactin production by *B. subtilis* on root surface during the process of colonization is also not known. Our data shows that mature region of the root is preferred by *B. subtilis* colonization during hydroponic culturing at 25°C. Contrary to the literature, we showed that *B. subtilis* irrespective of bacterial derived surfactin prefers to bind at the MR region of the roots. The mature region (MR) of the roots is an important rhizoplane frontier, this root zone initiates the formation of lateral primordia (29) and may secrete specific metabolites which may trigger *Bacillus* colonization. Interestingly, our data showed that microbially-derived surfactin may not suffice the root-microbe interaction initiation and colonization, and an exogenous level of surfactin may be needed to trigger colonization. This was shown, when an exogenous surfactin was added in the surfactin null strain (PY79), leading to root colonization in the MR region.

However, under abiotic conditions, we noted that sfp^+^ formed pellicles similar to that of the undomesticated strain UD1022, while PY79 was unable to form a complete and continuous pellicle. This aligns with the findings of Stoll et al. (1), wherein it was shown that the production of surfactin assists in the formation of biofilms and microcolony initiation. However, the production of surfactin or lack thereof was not the sole determinant of whether a microbe would be a successful colonizer or not (1). Other studies have also noted that even without surfactin, some *Bacillus* species are still able to form weak biofilms if they are able to produce NRP such as an iturin or bacillomycin (28). One of the links to surfactin’s role in biofilm production is the linkage to the activation of the gene *spo0A* which produces a response regulator protein (30). Even with a knockout of surfactin, many species were still able to produce a pellicle despite the changes in morphology and loss of the ability to float (30). A recent study showed that surfactin may be involved in triggering the rate of pellicle formation (31). The study shows that supplementation of surfactin may enhance the cell density in advancing pellicle timing (31). The study also indicates that surfactin biosynthesis may vary in different *B. subtilis* strains as the pellicle timings forms a critical factor in formation of biofilms on abiotic surfaces. In accordance, our data also showed that supplementation of surfactin did not alter pellicle formation in surfactin null strain PY79/sfp^-^. The surfactin overproducing strain sfp^+^ showed a continuous pellicle formation compared to the surfactin null strain, suggesting that microbial surfactin production may help in pellicle formation on abiotic surfaces. The ability of an undomesticated strain UD1022 to form pellicle was like sfp^+^ strain, suggesting that surfactin levels in both strains may be similar. Both sfp^+^ and undomesticated strains showed similar pellicle formation under all the treatments in two different incubation temperature (22°C and 30°C) conditions. This data suggests that surfactin-mediated pellicle formation is independent of the incubated temperature conditions in our studies. Most reports showing pellicle formation in the literature uses 30°C as the incubation temperature as a conducive condition for bacterial growth (32). The idea to test pellicle formation as a factor of surfactin production under a lower temperature was to mimic the root association studies with *B. subtilis* and plant hosts. The effect of surfactin and its role in both biotic and abiotic surface attachment/association in a gnotobiotic system either for root association or for pellicle formation are difficult to tease out the underlying mechanisms, thus, future experiments should include gene expression coupled with microscopy to assess the effect of specific root zones in triggering root attachment at a single cells level.

The amino acid glutamate (Glu) is an important signaling molecule in plants, playing roles in many important functions including wound response, pathogen resistance, response/adaptation to abiotic stress (33). Glutamate has been observed being released from the maize roots under duress from iron (Fe) deficiencies in the soil (34). In addition, glutamate has been shown to trigger a Ca^2+^ transient based systemic signaling response against herbivory (20). Glutamate also plays a role as a plant resistance inducer (PRI) activating plant defense against pathogenic fungi (35). However, studies regarding the impact of glutamate in the rhizosphere to initiate root association or signaling with PGPRs are lacking. The role of glutamate to either encourages or deters colonization of roots appears to be unknown. In our study we observed an increase in the root colonization when plants and bacteria were exposed to glutamate exogenously and when plants were primed with glutamate. The data clearly showed that supplementation of glutamate overrides surfactin for root association even in a surfactin null strain. The data showing the role of exogenous Glu on surfactin titers in static bacterial culture/plant associated sfp+ strain dictates that surfactin biosynthesis and Glu may be operating differently for bacterial colonization and are condition specific responses. PGPRs respond not only to biotic stressors like disease, but abiotic stressors as well (36). Since increased bacterial root attachment was observed when glutamate was added to the media, it could be possible that the microbes were responding to a perceived abiotic stressor occurring to the plant and following the signal to attach to help mitigate said stress. In accordance, Kim et al. (37) also indicated that internal levels glutamic acid of the plant is important the cultivation of beneficial microbial communities. The role of glutamate in triggering pellicle formation on the abiotic surface in both surfactin null and sfp+ strain as like non-glutamate treatments. The data shows that glutamate plays a critical role in triggering root association compared to its role in facilitating pellicle biofilm formation on an abiotic surface. The role of root surface chemistry and exudation from different root zones may be critical for bacteria attachment. It is known that benign microbes including *B. subtilis* induces innate root defense response (38). Benign microbes including *Bacillus* species have evolved to suppress plant root defense response to initiate colonization and biofilm formation. The role of surfactin and/or glutamate in suppressing root defense is not established. Arguably, it will be interesting to conceptualize that different root regions such as root tip, CEZ or MR may have a different root surface/secretion chemistry which may change the pattern of colonization by beneficial bacterial species (Concept Figure 8). Detailed studies are needed to decipher the role of glutamate (either plant derived, or soil-based) for initiation of attachment on both abiotic and biotic surfaces and for evaluating its role in modulating innate root defense response and secretion/surface chemistry.

**Figure 8:**
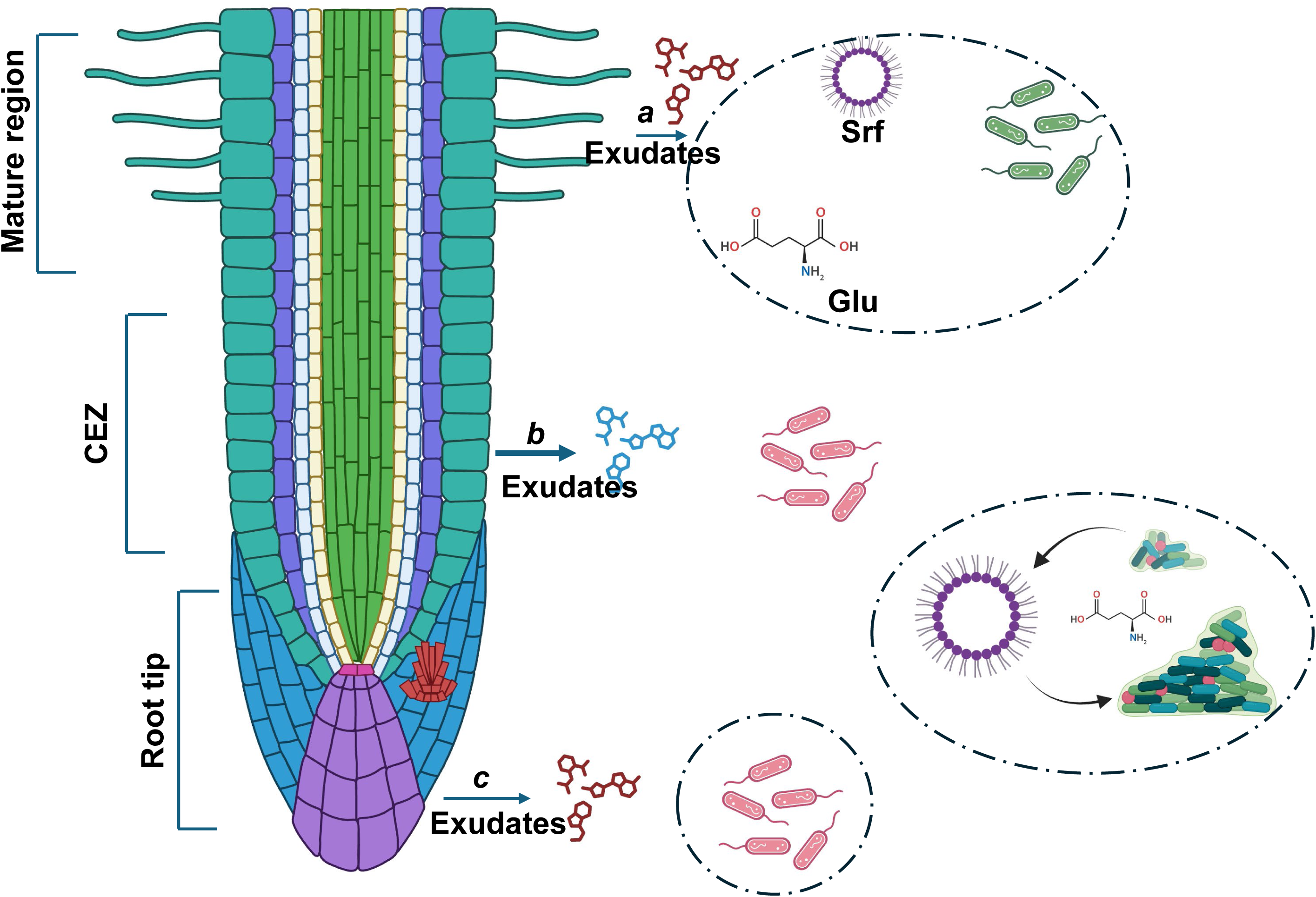
Schematic shows the root surface chemistry in the association of benign microbes. The root zones such as root tip and central elongation zone (CEZ) are speculated to be preferred zones of an association by benign microbes compared to the mature region (MR) of the roots. It is conceptualized that these three regions may have a different root surface chemistry which may change the pattern of colonization by beneficial bacterial species. Glutamate is shown to override innate defense response of plant roots for a robust colonization phenotype which is independent of surfactin production.

### Conclusion

PGPRs are vital for plant growth and health, however many of the mechanisms of how and where they attach to the plant root is largely unknown. To explore this largely unexplored area of science more thoroughly, we created detailed root maps of a young tomato plant. Through the creation of the root map, we were able to observe that most of the PGPR colonization appears to occur around the region of maturation (MR) of the root and not around the root tip or central zone of elongation. This trend could be due to specific exudates released from the plant around this area or due to the larger surface area as attachment was also observed around the root hairs. However, the presence or absence of microbially-derived surfactin did not seem to modulate root colonization. The addition of glutamate to the plant growth environment challenged the pattern of colonization, especially when plants were primed with glutamate prior to the addition of PGPRs. While microbes still colonized the mature region, in some cases they also colonized the root tip and central zone of elongation. The addition of surfactin to the media alone did not change these colonization patterns and only appeared to enhance colonization slightly. However, when glutamate was added into the media, colonization of the plant by of PY79 (surfactin null strain), increased dramatically. Abiotic surfaces of the pellicle experiments yielded different results from the biotic surfaces of the roots. Both an undomesticated strain UD1022 and a sfp+ formed pellicles under various conditions. UD1022 even managed to form wrinkles and continuous biofilm at 22℃. This differed from what was observed in experiments involving root surfaces with the PY79 treatment. While PY79 could not attach to root surfaces on its own, once glutamate or surfactin (or both) was used as a treatment, colonization could be observed. Under abiotic conditions, PY79 was unsuccessful in forming continuous pellicles in the treatments. It is fair to conclude that pellicle formation or lack thereof, may not be representative of the potential of a microbe to attach to biotic surfaces such as a root. Overall, surfactin production by PY79 didn’t impact root attachment in tomatoes, and other microbially-derived secondary metabolites should be investigated further. The role of surfactin to trigger root association may depend on the concentration of surfactin. In addition, the role of glutamate to trigger a surfactin independent root association opens areas to verify and test other PRI’s and amino acid derivatives to facilitate benign plant-microbe interactions. In addition, how glutamate modulates plant innate defense response and secretion chemistry to enhance root colonization needs to be studied at the mechanistic level.

## Supporting information

SOM Figures 1-3

## Acknowledgments

HPB and AMK acknowledges the support from USDA NIFA BRAG grant (2021-33522-35807). RW acknowledges the support from Microbiology Graduate Program at University of Delaware.

## Legends to the supplementary figure

**Supplementary Figure 1:** HPLC traces of surfactin analysis in sfp- and sfp+ strains treated with/without glutamate in a plant mesocosm suspension culture.

**Supplementary Figure 2: A-B:** The panel shows tomato root phenotypes with sfp- and sfp+ strains. Different bacterial densities (OD_600_= 0.01-0.0001) were inoculated with tomato plants (A), and plants post-treatment were recorded for root traits, such as length of primary roots (PR), length of lateral roots (LLR) and number of lateral roots (NLR) were quantified post 7 days of incubation (B). Letters indicate statistical similarity between the bacterial concentrations using one-way ANOVA and Tukey multiple comparisons (p-value < 0.05).

**Supplementary Figure 3: A-D:** The panel shows tomato root phenotype with sfp- and sfp+ treatment with/without glutamate (5mM) (A). Tomato plants were inoculated with sfp- and sfp+ strains (OD_600_=0.01). Plants post-treatment was recorded for root traits, such as length of primary roots (PR) (B), length of lateral roots (LLR) (C) and number of lateral roots (NLR) (D), which were quantified after 7 days of incubation. Two-way ANOVA was performed and Tukey multiple comparisons to determine statistical significance (p-value < 0.05). * = 0.0303, *** = 0.0008, **** = < 0.0001.

## References

1. Stoll A, Salvatierra-Martínez R, González M, Araya M. 2021. The Role of Surfactin Production by *Bacillus velezensis* on Colonization, Biofilm Formation on Tomato Root and Leaf Surfaces and Subsequent Protection (ISR) against *Botrytis cinerea*. Microorganisms 9(11):2251.

2. Banerjee S, van der Heijden MG. 2023. Soil microbiomes and one health. Nat Rev Microbiol.

3. Backer R, Rokem JS, Ilangumaran G, Lamont J, Praslickova D, Ricci E, Subramanian S, Smith DL. 2018. Plant growth-promoting rhizobacteria: context, mechanisms of action, and roadmap to commercialization of biostimulants for sustainable agriculture. Front Plant Sci 23;9:1473.

4. Vlamakis H, Chai Y, Beauregard P, Losick R, Kolter R. 2013. Sticking together: building a biofilm the Bacillus subtilis way. Nat Rev Microbiol 11(3):157–68.

5. Barreto, HC, Cordeiro, TN, Henriques, AO. Gordo I. 2020. Rampant loss of social traits during domestication of a *Bacillus subtilis* natural isolate. Sci Rep 10(1):18886

6. Gallegos-Monterrosa R, Mhatre E, Kovács ÁT. 2016. Specific Bacillus subtilis 168 variants form biofilms on nutrient-rich medium. Microbiology. 162(11):1922–32.

7. Luo C, Liu X, Zhou H, Wang X, Chen Z. 2015. Non-ribosomal peptide synthase gene clusters for lipopeptide biosynthesis in *Bacillus subtilis* 916 and their phenotypic functions. Appl Environ Microbiol. 81(1):422–31.

8. Ongena M, Jourdan E, Adam A, Paquot M, Brans A, Joris B, Arpigny JL, Thonart P. 2007. Surfactin and fengycin lipopeptides of Bacillus subtilis as elicitors of induced systemic resistance in plants. Environ Microbiol. 9(4):1084–90.

9. Rahman FB, Sarkar B, Moni R, Rahman MS. 2021. Molecular genetics of surfactin and its effects on different sub-populations of *Bacillus subtilis*. Biotechnology Reports, 32:e00686.

10. Hoff G, Arguelles Arias A, Boubsi F, Pršić J, Meyer T, Ibrahim HMM, Steels S, Luzuriaga P, Legras A, Franzil L, Lequart-Pillon M, Rayon C, Osorio V, de Pauw E, Lara Y, Deboever E, de Coninck B, Jacques P, Deleu M, Petit E, Van Wuytswinkel O, Ongena M. 2021. Surfactin stimulated by pectin molecular patterns and root exudates acts as a key driver of the *Bacillus*-plant mutualistic interaction. mBio. 1;12(6):e0177421.

11. Théatre A, Cano-Prieto C, Bartolini M, Laurin Y, Deleu M, Niehren J, Fida T, Gerbinet S, Alanjary M, Medema MH, Léonard A. 2021. The surfactin-like lipopeptides from *Bacillus* spp.: Natural biodiversity and synthetic biology for a broader application range. Frontiers in Bioengineering and Biotechnology 9:623701.

12. Bais HP, Weir TL, Perry LG, Gilroy S, Vivanco JM. 2006. The role of root exudates in rhizosphere interactions with plants and other organisms. Annu. Rev. Plant Biol. 57(1):233–66.

13. Beauregard PB, Chai Y, Vlamakis H, Losick R, Kolter R. 2013. *Bacillus subtilis* biofilm induction by plant polysaccharides. Proceedings of the National Academy of Sciences. 110(17):E1621–30.

14. Rudrappa T, Czymmek KJ, Paré PW, Bais HP. (2008) Root-secreted malic acid recruits beneficial soil bacteria. Plant physiology. 148(3):1547–56.

15. Killiny N, Hijaz F. (2016) Amino acids implicated in plant defense are higher in *Candidatus liberibacter asiaticus*-tolerant citrus varieties. Plant Signal Behav.;11: e1171449.

16. Popko M, Michalak I, Wilk R, Gramza M, Chojnacka K, Górecki H. (2018) Effect of the New Plant Growth Biostimulants Based on Amino Acids on Yield and Grain Quality of Winter Wheat. Molecules. 21 :470–478.

17. Kadotani N, Akagi A, Takatsuji H, Miwas T, Igarshi D. (2018) Exogenous proteinogenic amino acids induce systemic resistance in rice. BMC Plant Biol 16, 60.

18. Yang J, Sun C, Fu D, Yu T. (2017) Test for l-glutamate inhibition of growth of *Alternaria alternata* by inducing resistance in tomato fruit. Food Chem. 1;230:145–153.

19. Li ZG, Ye XY, Qiu XM. (2019) Glutamate signaling enhances the heat tolerance of maize seedlings by plant glutamate receptor-like channels-mediated calcium signaling. Protoplasma. 256(4):1165–1169.

20. Toyota M, Spencer D, Sawai-Toyota S, Jiaqi W, Zhang T, Koo AJ, Howe GA, Gilroy S. 2018. Glutamate triggers long-distance, calcium-based plant defense signaling. Science 361 (6407): 1112–15.

21. Hmidet N, Ben Ayed H, Jacques P, Nasri M. 2017. Enhancement of surfactin and fengycin production by *Bacillus mojavensis* A21: application for diesel biodegradation. BioMed research international. (1):5893123.

22. Liu S, Tang MH, Cheng JS. 2023. Fermentation optimization of surfactin production of *Bacillus amyloliquefaciens* HM618. Biotechnology and Applied Biochemistry. 70(1):38–50.

23. Earl AM, Losick R, Kolter R. 2008. Ecology and genomics of Bacillus subtilis. Trends Microbiol (*6*):269–75.

24. Pieterse CM, Zamioudis C, Berendsen RL, Weller DM, Van Wees SC, Bakker PA. 2014. Induced systemic resistance by beneficial microbes. Annual review of phytopathology, 52, 347–75.

25. Lakshmanan, V, Selvaraj G, Bais HP. 2014. Functional Soil Microbiome: Belowground Solutions to an Aboveground Problem. Plant Physiology 166 (2): 689–700.

26. Qi X, Liu W, He X, Du C. 2023. A review on surfactin: molecular regulation of biosynthesis. Arch Microbiol. 205(9): 313.

27. Liu Y, Xu Z, Chen L, Xun W, Shu X, Chen Y, Sun Y, Wang J, Ren Y, Shen Q, Zhang R. (2024) Root colonization by beneficial rhizobacteria, FEMS Microbiology Reviews, 48: 1 fuad066,

28. Thérien M, Kiesewalter HT, Auria E, Charron-Lamoureux V, Wibowo M, Maróti G, Kovács ÁT, Beauregard PB. 2020. Surfactin production is not essential for pellicle and root-associated biofilm development of Bacillus subtilis. Biofilm 2:100021.

29. Malamy JE. 2005. Intrinsic and environmental response pathways that regulate root system architecture. *Plant*, Cell & Environment 28 (1): 67–77.

30. Zhang Y, Qi J, Wang Y, Wen J, Zhao X, Qi G. 2022. Comparative study of the role of surfactin-triggered signalling in biofilm formation among different Bacillus species. Microbiological Research 254, 126920.

31. Stannius RO, Fusco S, Cowled MS, Kovács ÁT. (2024) Surfactin accelerates *Bacillus subtilis* pellicle biofilm development. Biofilm. 30; 9:100249.

32. Dragoš A, Kovács ÁT. 2017. The peculiar functions of the bacterial extracellular matrix. Trends in microbiology 25(4):257–66.

33. Qiu XM, Sun YY, Ye XY, Li ZG. 2020. Signaling role of glutamatetamate glutamate in plants.” Frontiers in Plant Science 10:1743.

34. Carvalhais LC, Dennis PG, Fedoseyenko D, Hajirezaei MR, Borriss R, Von Wirén N. 2011. Root exudation of sugars, amino acids, and organic acids by maize as affected by nitrogen, phosphorus, potassium, and iron deficiency. Journal of Plant Nutrition and Soil Science 174(1):3–11.

35. Goto Y, Maki N, Ichihashi Y, Kitazawa D, Igarashi D, Kadota Y, Shirasu K. 2020. Exogenous treatment with glutamate induces immune responses in Arabidopsis. Molecular Plant-Microbe Interactions. 33(3):474–87.

36. Kumar R, Swapnil P, Meena M, Selpair S, Yadav BG. 2022. Plant growth-promoting rhizobacteria (PGPR): Approaches to alleviate abiotic stresses for enhancement of growth and development of medicinal plants. Sustainability. 14(23):15514.

37. Kim DR, Jeon CW, Cho, G. et al. (2021) Glutamic acid reshapes the plant microbiota to protect plants against pathogens. Microbiome 9, 244.

38. Lakshmanan V, Kitto SL, Caplan JL, Hsueh YH, Kearns DB, Wu YS, Bais HP. 2012. Microbe-associated molecular patterns-triggered root responses mediate beneficial rhizobacterial recruitment in *Arabidopsis*. Plant Physiology. 160(3):1642–61.

