## Supplementary figures and images for "The influence of surfactin and glutamate on tomato root association by plant growth promoting rhizobacteria *Bacillus subtilis*"

### SOM Figures 1-3

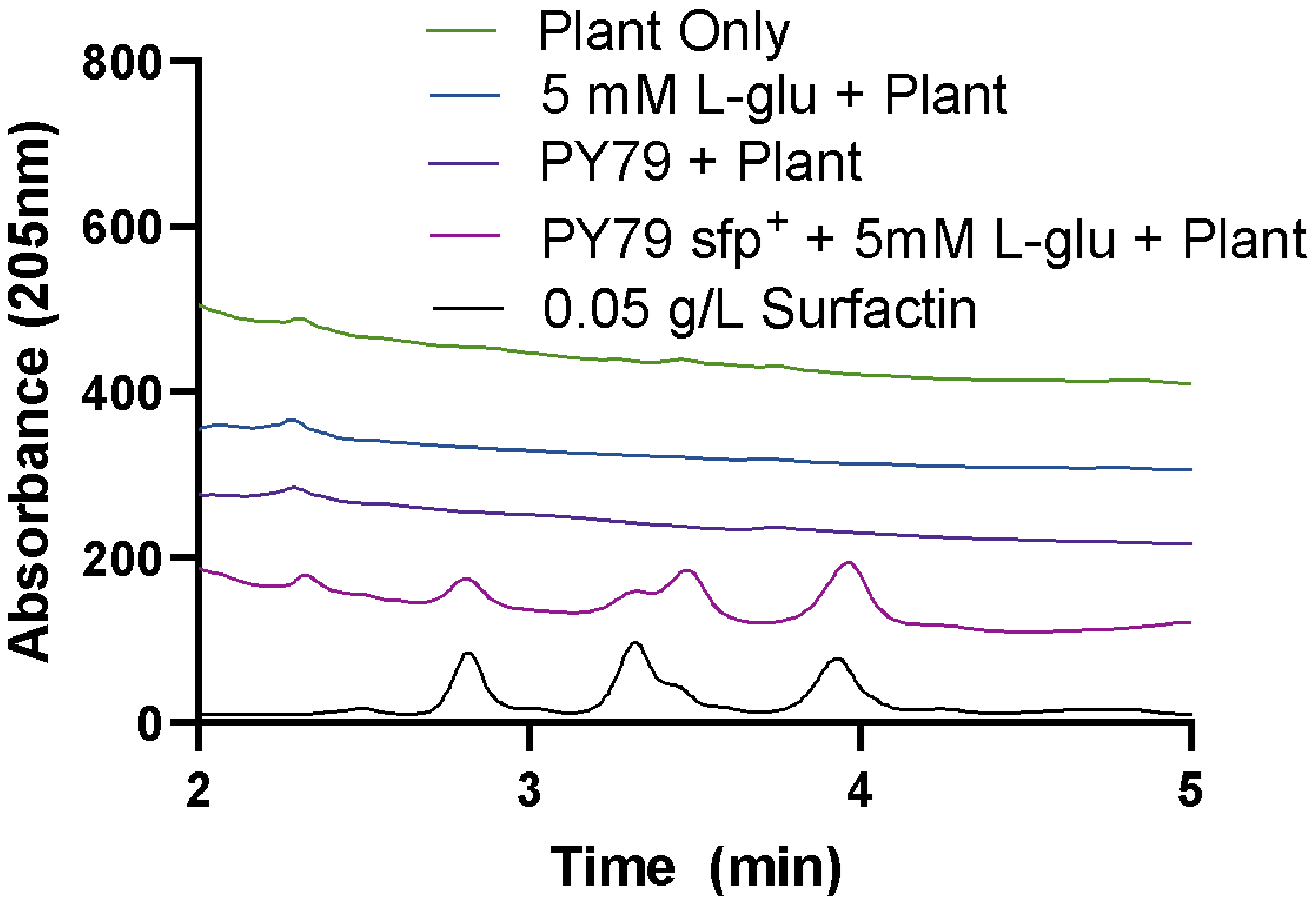

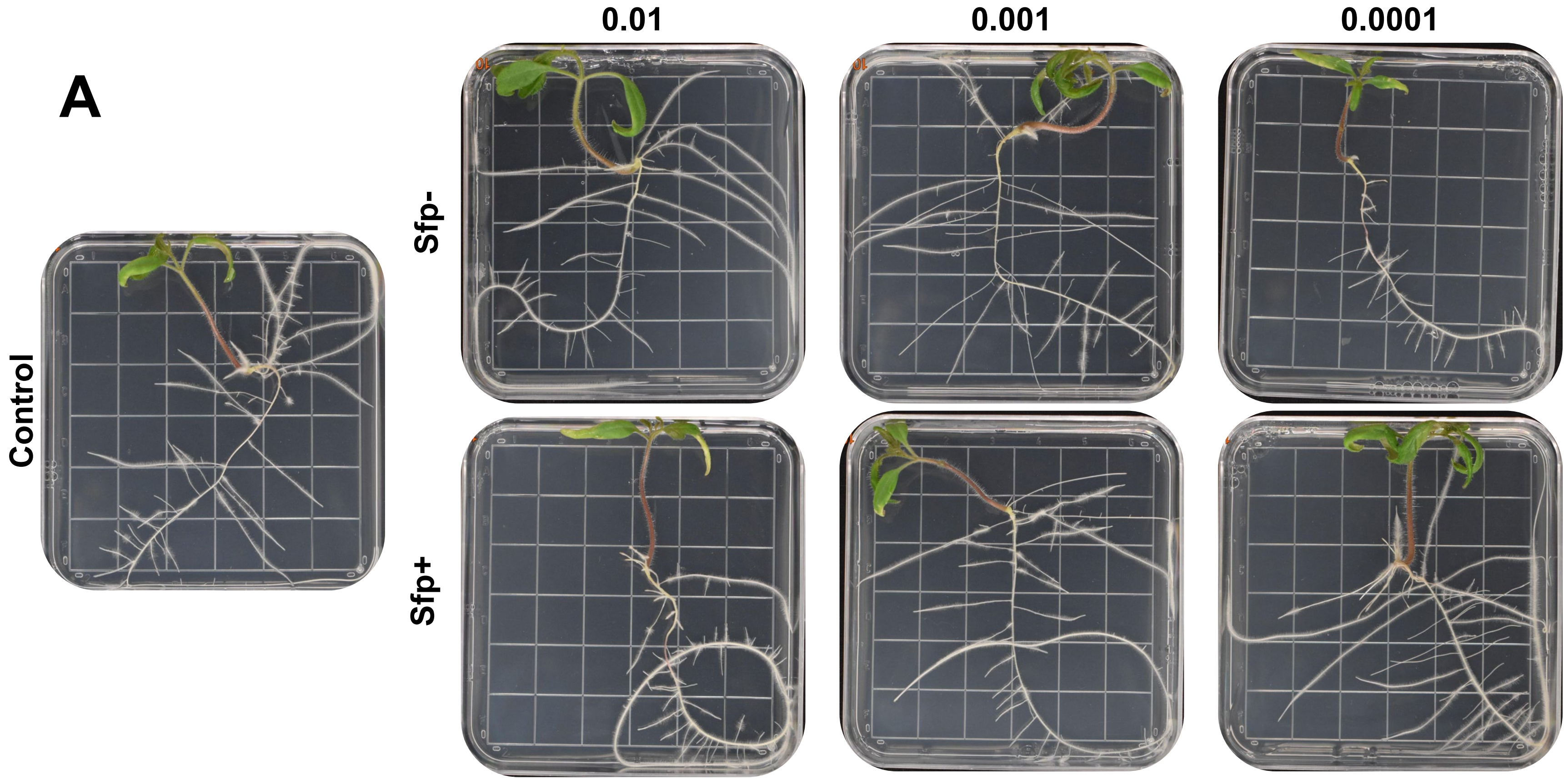

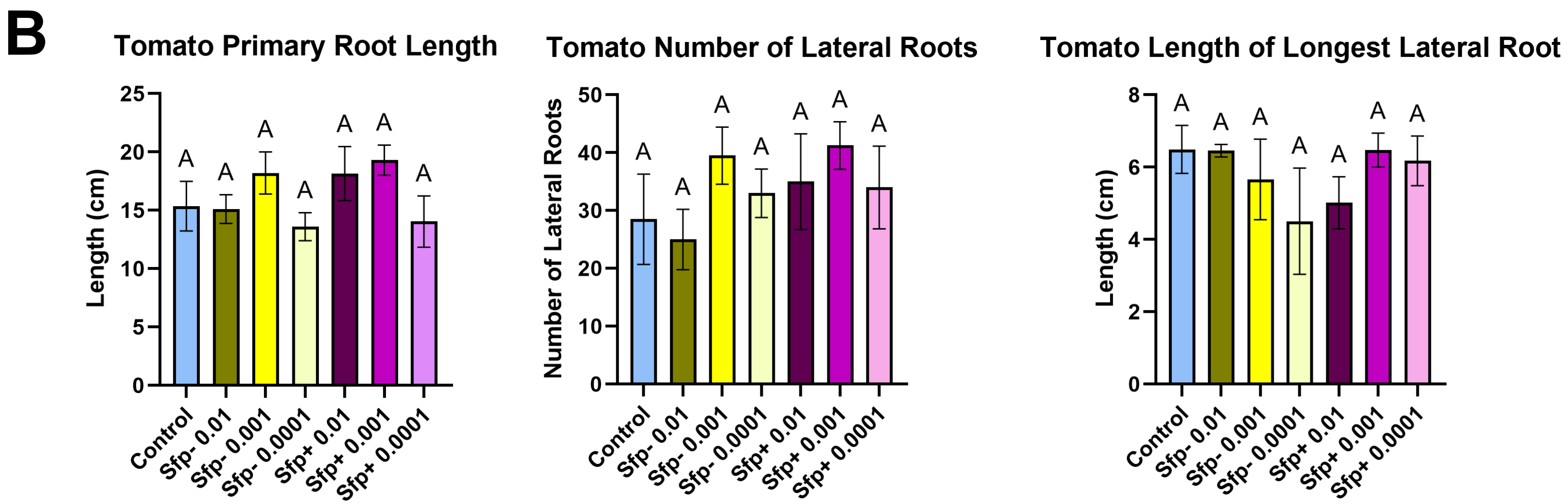

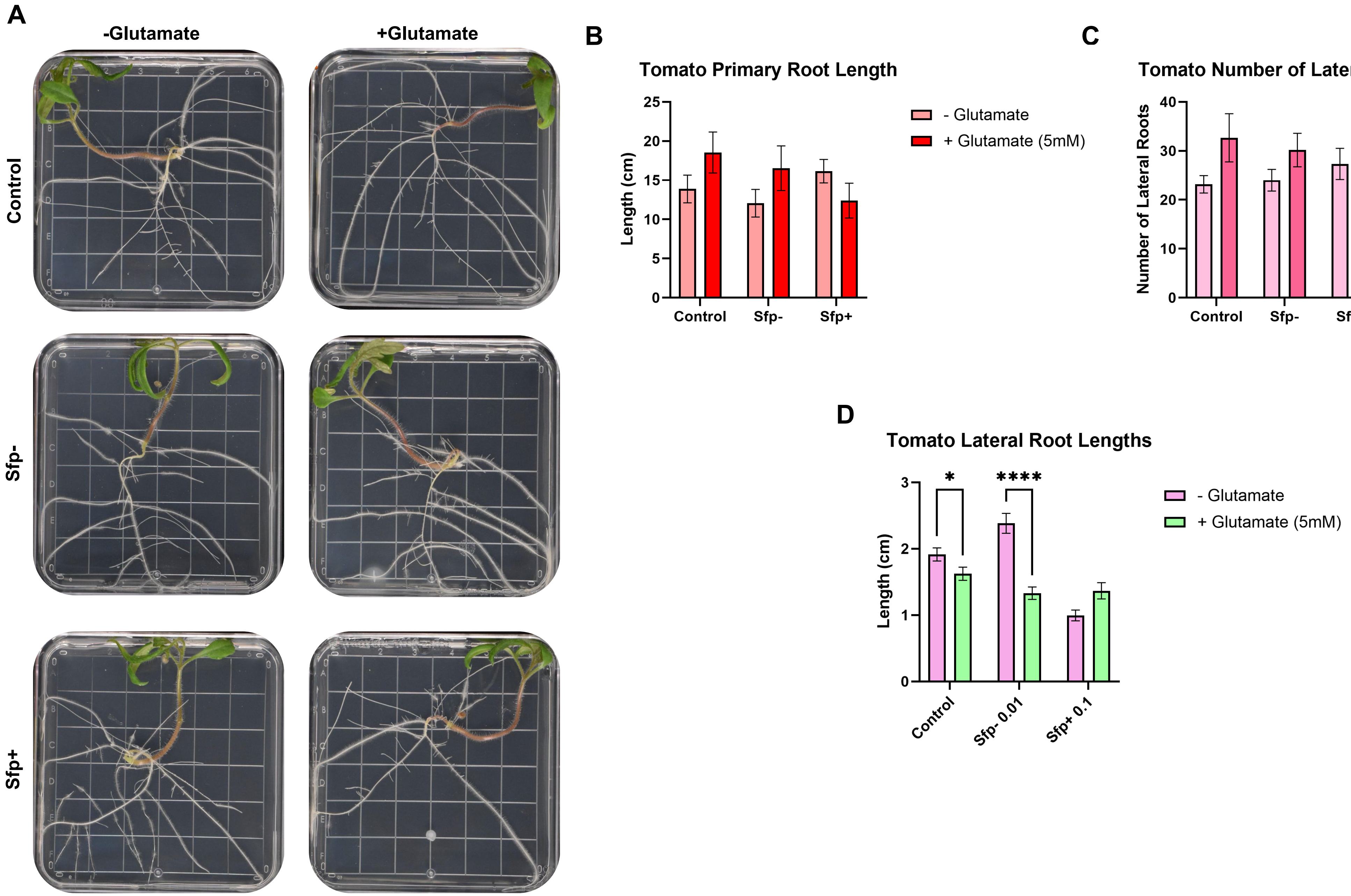
